# Human spinal cord activation during filling and emptying of the bladder

**DOI:** 10.1101/2024.02.16.580736

**Authors:** K. A. Agyeman, D.J. Lee, A. Abedi, S. Sakellaridi, E.I. Kreydin, J. Russin, Y.T. Lo, K. Wu, W. Choi, V.R. Edgerton, C. Liu, V.N. Christopoulos

## Abstract

Recording neural activity from the spinal cord is crucial for gaining insights into how it functions. However, the neural activity of the human spinal cord is notoriously difficult to measure. The bony and fascial enclosures combined with the relatively small anatomic size of the spinal cord make it an unfavorable target for traditional functional neuroimaging techniques. Functional ultrasound imaging (fUSI) is an emerging neuroimaging technology that represents a new platform for studying large-scale neural dynamics with high sensitivity, spatial coverage and spatiotemporal resolution. Although it was originally developed for studying brain function, fUSI was recently extended for imaging the spinal cord in animals and humans. While these studies are significant, their primary focus is on the neuroactivation of the spinal cord in response to external sensory stimulations. Here, we combined fUSI with urodynamically-controlled bladder filling and emptying to characterize the hemodynamic response of the human spinal cord during the micturition cycle. Our findings provide the first practical evidence of the existence of bladder pressure-responsive regions, whose hemodynamic signal is strongly correlated with the bladder pressure.

## Introduction

The spinal cord has been frequently neglected in the study of neural function. As a result, its anatomy and physiology are not as well understood as those of the brain. Yet, it represents the first evolutionary step in central nervous system development and houses the neural circuitry that controls and modulates some of the most important functions of life ^1^. Neural networks capable of producing autonomous central commands – usually stereotyped and rhythmic motor behaviors – are present throughout the rostral and the caudal parts of the spinal cord^2^. Actions such as chewing, swallowing and breathing are thought to be partially produced by these networks in the rostral cord ^3^. Similarly, autonomic functions such as urination and defecation are under control of neural networks located in the caudal spinal cord ^4^.

Although evidence for the existence of neural network circuits that control and regulate certain body processes is strong, its demonstration in humans has been challenging to achieve. The bony, fascial enclosure and small cross-section dimensions (approximately 12 mm in diameter) of the spinal cord combined with susceptibility artifacts due to local magnetic field inhomogeneities generated by interfaces between surrounding bones, ligaments, soft tissues and cerebrospinal fluid (CSF) make the spinal cord an unfavorable target for traditional neuroimaging techniques, such as functional magnetic resonance imaging (fMRI) ^5–11^. As a result, the bulk of our understanding of spinal cord function comes from animal and lesioning studies ^12^. There is little direct evidence for function-specific spinal cord activity in humans, and fMRI – which has shed so much light on brain functions in humans – of the spinal cord is only minimally developed and generally restricted to the cervical cord^8–10,13^. Given this context, there is a clear and distinct need for developing neurotechnologies that make the functional study of the human spinal cord more accessible.

Functional ultrasound imaging (fUSI) is an emerging neuroimaging technology that represents a new platform with high sensitivity, spatial coverage and spatiotemporal resolution, enabling a range of new pre-clinical and clinical applications^14–23^. It was originally developed for brain neuroimaging in small animals (i.e., rodents)^16^. Based on power Doppler imaging, fUSI measures changes in cerebral blood volume (CBV) by detecting backscattered echoes from red blood cells moving within its field of view^24,25^. While fUSI is a hemodynamic technique, its superior spatiotemporal performance (i.e., 100 μm and up to 10 ms) and sensitivity (∼ 1 mm/s velocity of blood flow) offer substantially closer connection to the underlying neuronal signal than achievable with other hemodynamic methods such as fMRI. It is minimally invasive and requires a trephination in large organisms to enable the penetration of the ultrasound waves, as the skull attenuates the acoustic wave. The fUSI scanner is like any clinical ultrasound machine, making the unit freely mobile between different settings and negates the need for extensive infrastructure inherent to fMRI.

Recently, fUSI was extended to study the spinal cord responses to electrical and mechanical stimulations in small animals and human patients ^26–30^. Despite the significant contribution of these studies in understanding how the spinal cord reacts to external sensory stimulations, none of them have demonstrated spinal cord circuits associated with physiological functions (i.e., body processes) in humans. In the current study, we utilize fUSI to study the hemodynamic response of the spinal cord during urinary bladder filling and emptying in patients, undergoing general anesthesia and epidural spinal stimulation surgery for chronic low back pain treatment. By combining fUSI recordings from the spinal cord with intravesical bladder pressure recordings, we identified spinal cord regions in which the hemodynamic signal is strongly correlated with bladder pressure. Overall, our study provides the first in-human application of fUSI to characterize the hemodynamic response of the spinal cord during urodynamically-controlled bladder filling and emptying, opening new avenues for better understanding the mechanisms of control that the spinal cord exerts over micturition.

## Results

To investigate how human spinal cord hemodynamics respond to bladder filling and emptying process, we acquired fUSI images of the spinal cord from four (4) chronic low back pain patients, who underwent standard-of-care implantation of an epidural spinal cord stimulation (ESCS) device under general anesthesia (Fig. 1A). Note that the urodynamic experiment was performed before ESCS implantation. A miniaturized 15.6-MHz, 128-channel, linear ultrasound transducer array was inserted through a partial laminar opening onto the dura at the level of the 10th thoracic vertebra (T10) with a transverse field of view (Fig. 1A). We utilized a protocol that consisted of about 26 min of continuous fUSI signal acquisition, including 5 min of baseline activity, followed by 2 bladder filling cycles and 1 emptying cycle, interspersed by 2 hold periods (about 1 min each) (Fig. 1B). The bladder was filled and emptied, accompanied by continuous intravesical bladder pressure recordings using a Laborie Goby™ (Vermont, USA) urodynamics system. The same protocol was employed for all patients. Fig. 1C depicts the changes of the bladder pressure during filling and emptying for all 4 patients.

**Figure 1.**
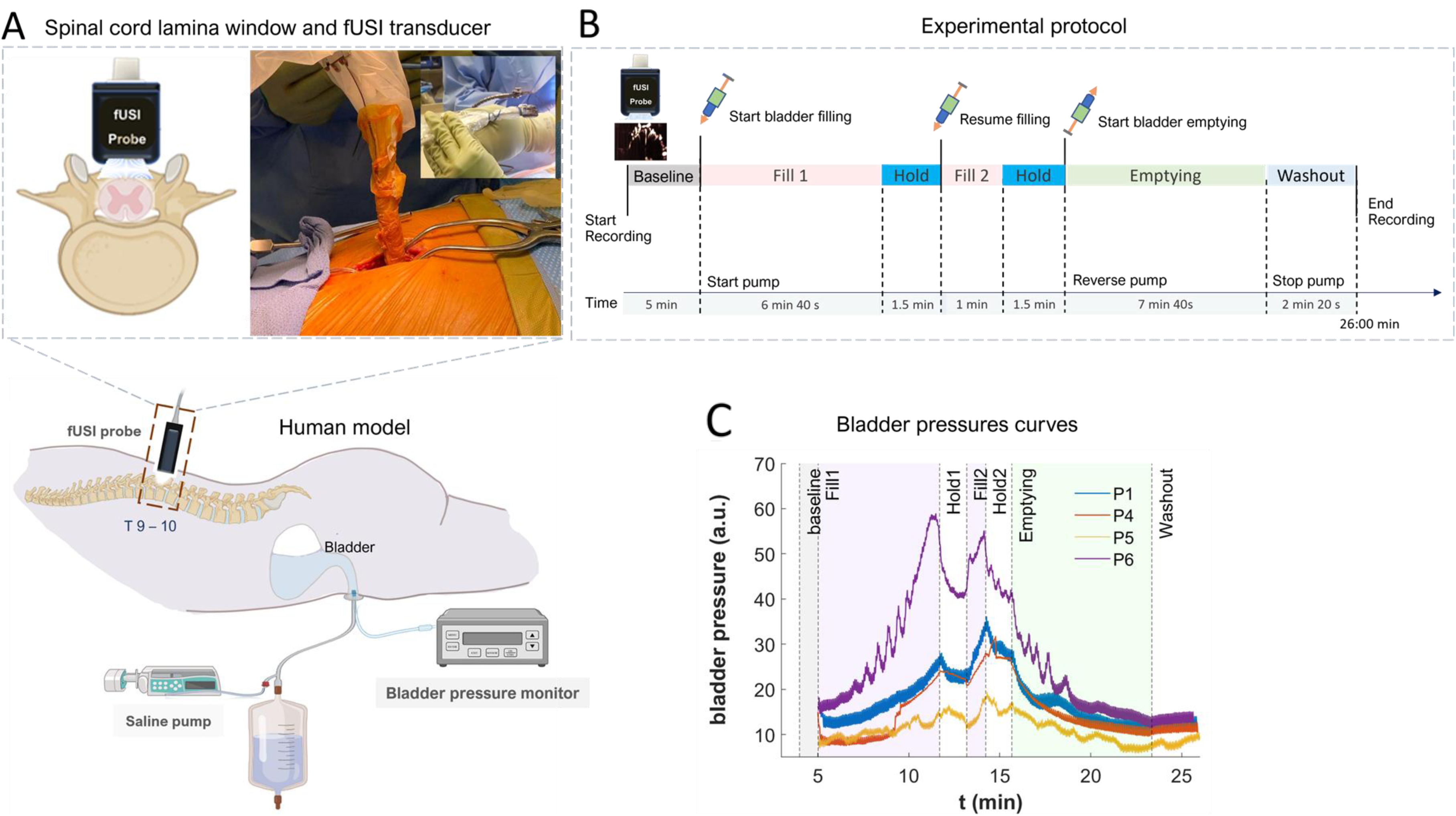
Experimental setup and fUSI acquisition protocol. **A)** A graphical representation of the human urodynamic model developed to study how the spinal cord activity is correlated with the bladder pressure. The spinal cord fUSI acquisition performed through a laminar window using a miniaturized 15.6-MHz, 128-channel, linear ultrasound transducer array. **B**) The experimental protocol for urodynamically-controlled filling and emptying the bladder. (**C**) Bladder pressure recordings across time during filling and emptying the bladder for the 4 patients.

### Hemodynamic response induced by bladder filling and emptying

Power Doppler (pD)-based functional ultrasound images were acquired from the spinal cord (Fig. 2). We used the mean spinal cord pD signal (1 min just before filling onset) to capture the anatomical vascularization of the human spinal cord in all patients, with the dorsal surface indicated by the white discontinuous lines (Fig. 2B). The pD images have spatial resolutions of 100 μm × 100 μm in-plane, plane thickness of about 400 μm, and a large field of view (FOV) 12.8 mm × 10 mm. The FOV captures the dorsal and portions of the ventral cross-section of the spinal cord – approximately indicated by the light-green rectangular overlay in Fig. 2A.

**Figure 2.**
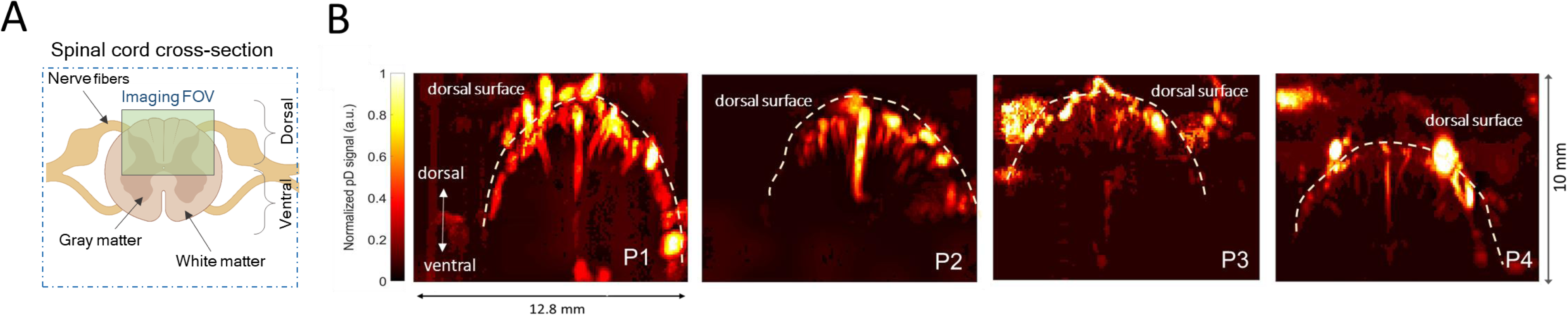
Functional ultrasound imaging of the spinal cord in a transverse plane. **A**) Cross section of spinal cord anatomy. The green area illustrates approximately the field of view of fUSI acquisition. (**B**) Power Doppler-based vascular maps showing the transverse section of the spinal cord of the four patients.

To characterize the spinal cord hemodynamic response during filling and emptying of the bladder, we computed the spinal cord blood volume changes (ΔSCBV) – i.e., % pD signal changes – relative to the baseline activity (i.e., average fUSI activity 1 min prior to start filling the bladder). The goal was to identify regions within the spinal cord that are correlated with bladder pressure. To do so, we computed the *activation map* for each patient by performing a Pearson’s correlation analysis between the bladder pressure changes and ΔSCBV for each pixel in the recorded area. The activation maps revealed spinal cord regions that are positively (reddish areas, r > 0.35, p < 0.01) and negatively (blueish areas, r <-0.35, p < 0.01) correlated with bladder pressure during filling and emptying the bladder (Fig. 3A). Notably, we observed bladder pressure-related regions extending beyond the dorsal surface, indicating that neural signals associated with bladder function may modulate hemodynamic activity in regions adjacent to the gray matter of the spinal cord. It is also likely that the activation detected in vessels outside the dorsal column may be attributed to their role in supplying blood to the vasculature within the gray matter.

**Figure 3.**
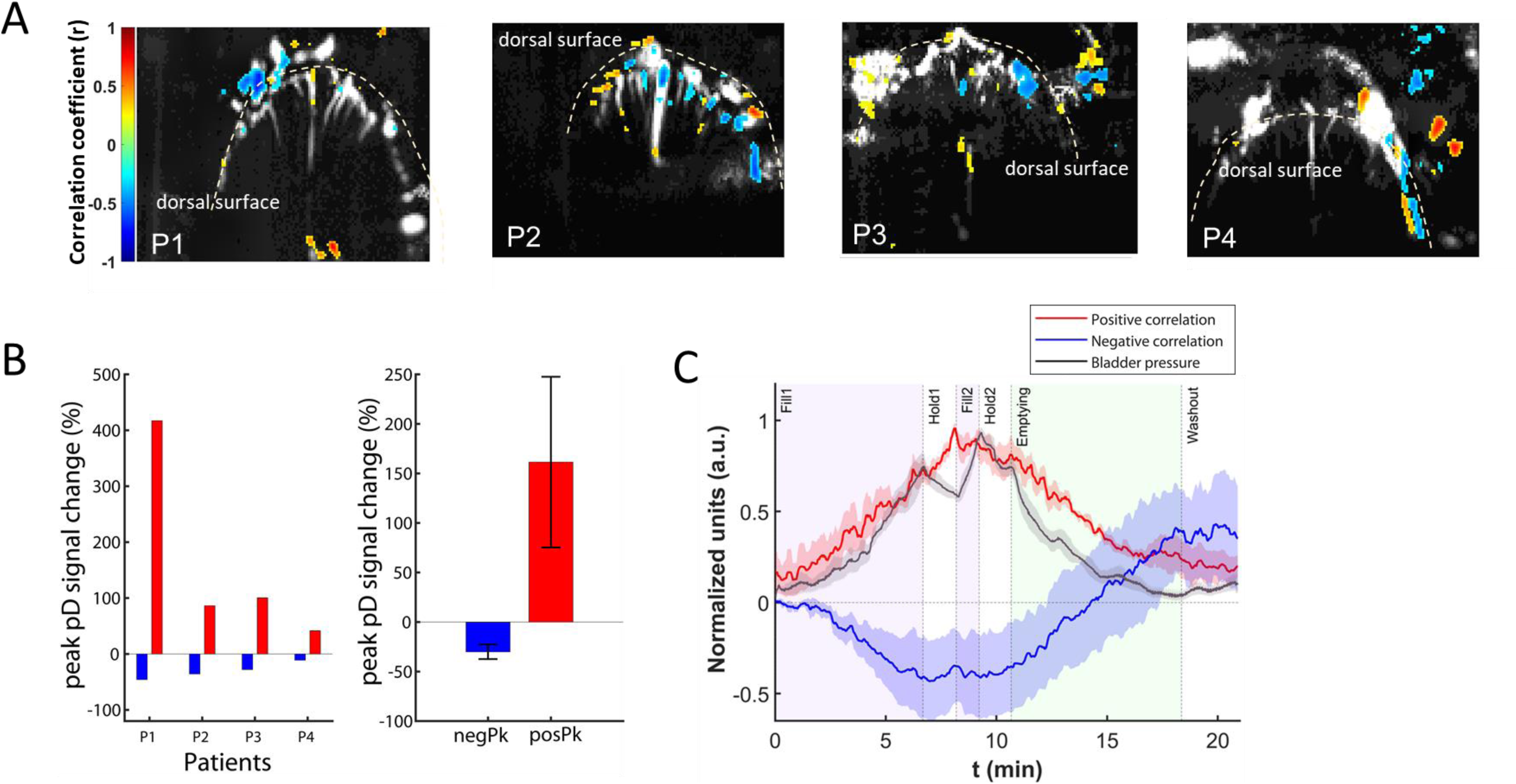
Activation maps of the correlation between ΔSCBV and bladder pressure during filling and emptying the bladder. **A**) Activation maps of the 4 patients that illustrate spinal cord regions that are positively (reddish) and negatively (blueish) correlated with the bladder pressure during filling and emptying the bladder. **B**) *Left panel*: Average ΔSCBV (i.e., % pD signal changes) from the baseline activity of bladder pressure-related regions for each of the 4 patients. Positive correlations with bladder pressure are depicted in red, while negative correlations are shown in blue. *Right panel*: Same as the left panel but across all 4 patients. **C**) Average normalized ΔSCBV of the spinal cord regions that are positively (red curve) and negatively (blue curve) correlated with the bladder pressure across patients. The gray curve depicts the normalized changes of the bladder pressure during the urodynamic experiment. The shaded regions around the bladder pressure and the ΔSCBV curves represent the standard error derived from averaging across patients.

To assess the temporal pattern of activation of the bladder pressure-related regions, we computed the average ΔSCBV over the pixels of the positive and negative correlates to the bladder pressure, across time and patients. Since the magnitude of the hemodynamic response changes varies between patients (Fig. 3B), we normalized the ΔSCBV between [-1, 1]. Similarly, we normalized the bladder pressure between [0, 1] to account for the different magnitudes of the pressure curves across patients. The results presented in Fig. 3C showed that bladder filling and emptying cause strong neuroactivation in the spinal cord. The regions positively correlated with the bladder pressure signal (i.e., red regions) exhibited a gradual increase in ΔSCBV during bladder filling with a subsequent gradual decrease in ΔSCBV during bladder emptying. Conversely, the regions negatively correlated with bladder pressure (i.e., blue regions) exhibited the opposite behavior – i.e., gradual decrease followed by increase of ΔSCBV during filling and emptying of the bladder, respectively. The gray curve depicts the average normalized bladder pressure changes across patients. The shaded regions around the bladder pressure and the ΔSCBV curves represent the standard error of mean across patients. The correlation between the bladder pressure and ΔSCBV is 0.89 ± 0.02 (Mean ± SE) for the positively (reddish) and -0.78 ± 0.05 for the negatively (blueish) bladder pressure-related spinal cord regions across patients.

### A machine learning algorithm to identify bladder pressure-related regions

We investigated whether we could detect spinal cord regions that encode the bladder pressure dynamics without directly monitoring the bladder pressure. To do so, we developed a machine learning technique to identify bladder pressure-related regions in the recorded images (Fig. 4). After collecting the fUSI data from the human spinal cord (Fig. 4A), we implemented a class-wise principal component analysis (cPCA) to reduce the dimensionality of the spinal cord imaging data (91×128 pixels per acquired image), and extracted effective discriminant features to differentiate between bladder filling (class:0, c0) and emptying (class:1, c1) classes (Fig. 4B). The entire fUSI spinal cord time series data acquired during filling and emptying periods were utilized (the hold time periods were excluded). The analysis was performed separately for each patient in whom the bladder pressure was successfully recorded (N=4). cPCA has been used to reduce sparsity and dimensionality while maintaining enough components to retain over 95% variance in the data (see Materials and Methods section for more details). It is ideally suited for discrimination problems with large dimensions and small sample size including natural and biomedical images ^31,32^. We paired cPCA with a class-discriminant support vector machine (SVM) classifier to determine the best decision boundary that separate the two classes - i.e., filling vs. emptying (Fig. 4C). A subset from each class was then separated into training (80%) and testing (20%) sets for cross-validation analysis. This approach results in a 1D low-dimension subspace that represents a feature extraction mapping from the 2D spinal cord image space. The subspace identifies pixels in the spinal cord fUSI images that encode differences between the filling (c0) and emptying (c1) classes, when projected back to the image space. Each pixel was assigned a relative weight of relevance normalized between [-1 1] – pixels with values close to +1 or -1 imply important components, while pixels with values close to 0 are less important with their fluctuations likely due to noise with respect to each class (Fig. 4C). Physically, the weighted regions can be interpreted as spinal cord regions in where ΔSCBV encodes differences between the filling and emptying classes. The positive and negative weights indicate that ΔSCBV contributes positively and negatively to the variation captured by the principal component, respectively. Hence, ΔSCBV of pixels that have positive and negative relative weights are positively and negatively correlated with the bladder pressure. The heat-map in Fig. 4D (left-top column) represents a typical example of the decoding analysis that identifies the most relevant spinal cord image pixels associated with filling and emptying the bladder for patient 1 – the most positive (reddish) and negative (bluish) relative weighted pixels overlaid on a grayscale mean fUSI spinal cord vascular map. Fig. 5A depicts the top 5% of the most heavily weighted pixels generated by the cPCA+SVM algorithm. The results showed that the average ΔSCBV of the activated regions was correlated with the bladder pressure with r = 0.65 ± 0.09, (mean ± SE) for the positive weights and r = -0.62 ± 0.08 for the negative weights across the 4 patients (Fig. 5C). Notably, the cPCA+SVM algorithm identified bladder pressure-related regions with less variability on the magnitude of the hemodynamic changes (i.e., %pD signal changes) during filling/emptying the bladder (Fig. 5B) compared to the original Pearson’s correlation analysis and the activation maps (see Fig. 3B).

**Figure 4.**
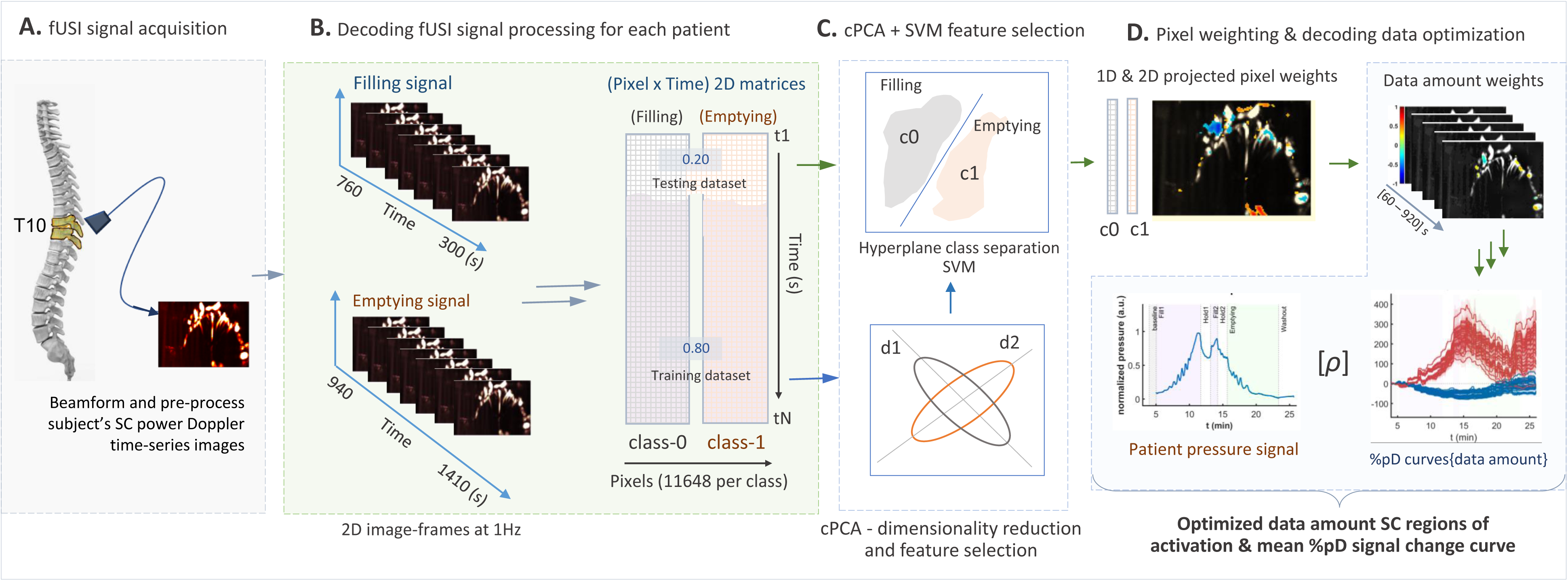
Flowchart of the cPCA+SVM algorithm developed to detect bladder pressure-related spinal cord regions. **A)** fUSI data during filling (class 0, c0) and emptying (class 1, c1) the bladder were recorded at the level of the T10 vertebral body and **B**) separated in training images and testing images based on the cross-validation technique used – 80% training data and 20% testing data. **C**) cPCA was paired with SVM to classify the state of the bladder (i.e., class 0 vs. class 1) using only the recorded pD signal from the spinal cord. **D**) This approach results in a 1-dimensional subspace the represents a feature extraction mapping from the 2D spinal cord image space. The subspace identifies pixels that are assigned with a relative weight between [-1, 1] and encodes the differences between the two classes – the higher the weight, the more significant the contribution of this pixel to the class separation. Highlighted panel shows the process for identifying the optimal amount of fUSI data that generate the best correlate between ΔSCBV and bladder pressure.

**Figure 5.**
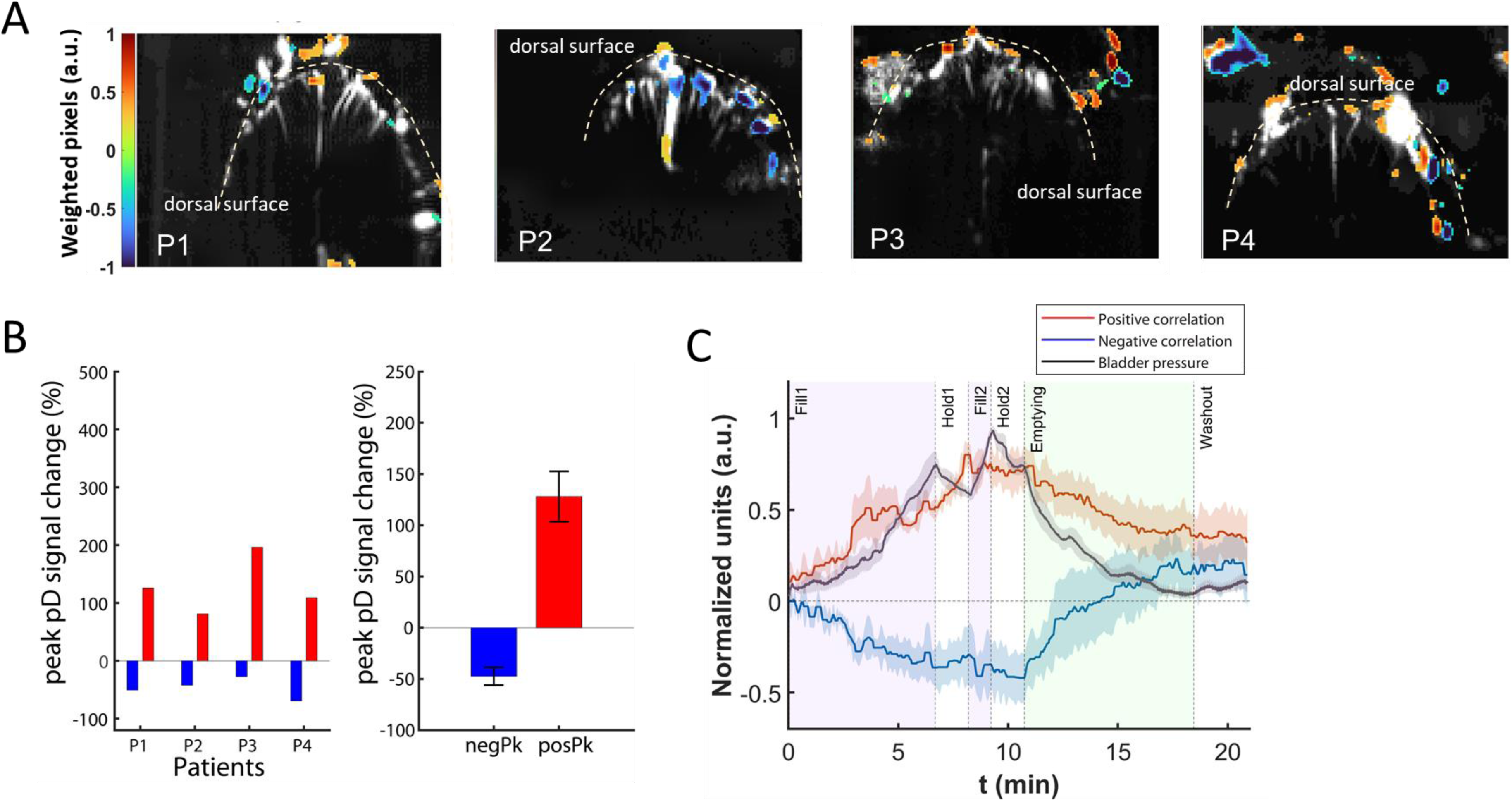
Bladder pressure-related spinal cord regions identified using cPCA+SVM. **A)** Weighted map of patients P1 to P4 extracted by the cPCA+SVM algorithm using all fUSI recorded data. The top 5% most heavily weighted voxels are shown. **B**) *Left panel*: Average ΔSCBV from the baseline activity of bladder pressure-related regions for each of the 4 patients. Positive correlations with bladder pressure are depicted in red, while negative correlations are shown in blue. *Right panel*: Same as the left panel but across all 4 patients. **C**) Average normalized ΔSCBV of spinal cord regions with positive weights (red curve) and negative weights (blue curve) as extracted by the cPCA+SVM algorithm using all fUSI data across the 4 patients. The shaded regions around the bladder pressure and the ΔSCBV curves represent the standard error derived from averaging across patients. Note that positive and negative weights correspond to positive and negative correlations of ΔSCBV with the bladder pressure.

### Optimal amount of data for decoding bladder pressure dynamics

So far, we have demonstrated that cPCA+SVM algorithm can accurately identify bladder pressure-related spinal cord regions. An interesting question is whether we can improve the cPCA+SVM performance using a subset of fUSI data – instead of entire data set – to train the classifier. The goal is to determine the optimal amount of data needed to train the classifier to detect the spinal cord regions that produce the best performance – i.e., the highest correlation between ΔSCBV of the bladder pressure-related spinal cord regions (i.e., as extracted by the cPCA+SVM algorithm) and the bladder pressure dynamics. To do so, we started with the last 30 s of fUSI data acquired during bladder filling for class c1, and the first 30 s of fUSI data acquired during bladder emptying for class c0. We employed 10-s data increment for each class – i.e., positive increment for class c0, and negative increment for class c1 – to derive a cumulative set of fUSI images that was used to train and evaluate the performance of the algorithm. We utilized similar cPCA+SVM decoding steps as outlined above with increasing data amounts. Each subset of data produced weighted relevant pixels that best discriminate between the two classes. A characteristic example of the weighted relevant pixels with the corresponding average ΔSCBV time course curves is illustrated in Fig. 4D (right column), in which the red and blue curves represent the average ΔSCBVs in areas with positive and negative weights, respectively. We then determined the optimal amount of fUSI data for each patient, in which the ΔSCBV of the weighted pixels exhibit the highest correlation with the bladder pressure (Fig. 4D highlighted area). The results showed that the cPCA+SVM algorithm produced activation maps comparable to those generated by using all recorded fUSI data (Fig. 6A), yet with enhanced performance. Specifically, there was a greater correlation between ΔSCBV and bladder pressure when utilizing a subset of the recorded fUSI data, as opposed to the entire dataset – i.e., r = 0.81 ± 0.05, (mean ± SE) for the positive weights when using 4.89 ± 0.57 min of the recorded fUSI images, and r = -0.85 ± 0.03 for the negative weights when using 3.58 ± 1.28 min of the recorded fUSI images, across the 4 patients (Fig. 6C). Notably, the variability of the %pD signal change during filling/emptying the bladder across the 4 patients was comparable to those generated when utilizing all the amount of recorded fUSI data (Fig. 6B).

**Figure 6.**
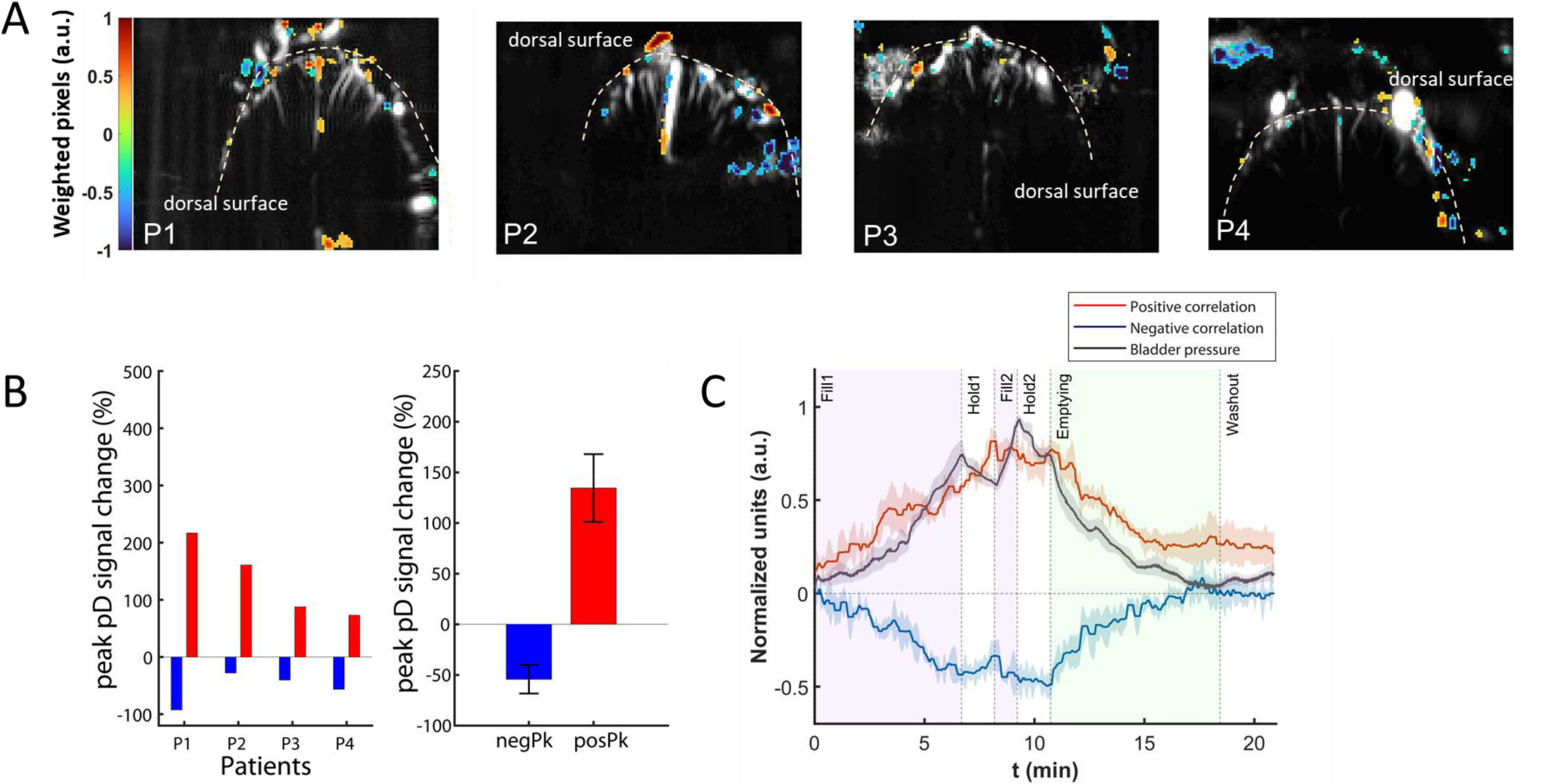
Bladder pressure-related spinal cord regions identified using cPCA+SVM with an optimal subset of fUSI recorded data. **A)** Weighted map of patients P1 to P4 extracted by the cPCA+SVM algorithm using an optimal subset of fUSI recorded data. The top 5% most heavily weighted voxels are shown. **B**) *Left panel*: Average ΔSCBV from the baseline activity of bladder pressure-related regions for each of the 4 patients. Positive correlations with bladder pressure are depicted in red, while negative correlations are shown in blue. *Right panel*: Same as the left panel but across all 4 patients. **C**) Average normalized ΔSCBV of spinal cord regions with positive weights (red curve) and negative weights (blue curve) as extracted by the cPCA+SVM algorithm using an optimal subset fUSI data across the 4 patients. The shaded regions around the bladder pressure and the ΔSCBV curves represent the standard error derived from averaging across patients. Note that positive and negative weights correspond to positive and negative correlations of ΔSCBV with the bladder pressure.

## Discussion

### General

While functional neuroimaging in the human brain has led to some progress in understanding brain function in micturition ^33–35^, the neural mechanism in the human spinal cord that controls filling and emptying of the bladder is almost entirely unclear. To the best of our knowledge, there is no study that has attempted to characterize hemodynamic changes in the spinal cord during filling and emptying of the bladder. One of the main reasons seems to be the intricate structure of the spinal cord, including its small cross-sectional area, the cardiac-related motion of cerebrospinal fluid (CSF), and motion artifacts caused by the proximity of organs such as the lungs. These factors make the spinal cord an unfavorable area for conventional functional neuroimaging studies ^5,8^. On the other hand, electrophysiology suffers from the inherent trade-offs between sampling density, coverage and channel count, making it challenging to achieve a spatial sampling resolution of less than 100 μm over a large recorded volume (i.e., 1 cm^3^ would require about 10^6^ channels). Optical imaging is capable of monitoring single-neuron activity over large areas, but is typically limited by a penetration depth of < 1 mm ^36,37^.

Within this context, fUSI represents an emerging neuroimaging technology that utilizes ultrasound to monitor blood flow changes as an indirect readout of neuronal activity with high spatiotemporal resolution, penetration depth and sensitivity to slow blood flow motion. Originally developed for brain neuroimaging, fUSI has been recently expanded to study spinal neurovascular responses in small animals ^26–29^ and human patients ^30^. Although these studies provide significant insights into better understanding the physiology of the spinal cord in sensory integrations, they are limited to artificial external stimulations, illustrating that fUSI is capable of detecting binary discrete spinal cord states– i.e., stimulation on vs. stimulation off. In the current study, we took the next major leap in fUSI spinal cord research by recording functional activity of the human spinal cord during urodynamically-controlled bladder filling and emptying. We showed that fUSI can detect spinal cord regions in which the hemodynamic signal is highly correlated with the bladder pressure. We also introduced a machine learning algorithm that can detect bladder pressure-related spinal cord regions, even when information about the bladder pressure is not available. Overall, our success in characterizing and correlating spinal cord hemodynamics to urodynamically-controlled micturition events holds promise for further understanding the functional and dysfunctional anatomy associated with lower urinary tract physiology.

### Neuroscience and scientific applications

The unique combination of fUSI technology with anatomically correlated and easily monitored physiological function of micturition – mimicked by urodynamically-controlled filling and emptying of the bladder – open new opportunities for better understanding of the spinal cord networks that promote urinary storage and induce urinary emptying. It also creates avenues for studying the neural circuitries that control and modulate other important bodily functions, such as sensation, ambulation (e.g., passive movements in anesthetized patients). Additionally, the existence of spinal cord regions, where the hemodynamic signals are strongly correlated with bladder pressure, provides the proof-of-concept for developing ultrasound-based spinal cord machine interface technologies for bladder control in patients with neurogenic bladder. Surveys have repeatedly revealed that restoration of bladder function remains the top priority for spinal cord injury patients, far ahead of even restoring the ability to walk^38^. In addition to spinal cord injury, a far greater number of people worldwide suffer from urinary dysfunctions of neurological origin. Developing spinal cord machine interfaces for informing the patients about the state of the bladder would be a step closer to restoring bladder control.

### New avenues for improving neuromodulation treatments for neurogenic lower urinary tract dysfunction

Urinary dysfunctions of neurological origin due to spinal cord or brain injury, degeneration, or stroke represent some of the biggest medical burdens in the world and lead to uniquely dehumanizing consequences ^39^. Therapies that are currently available abate some symptoms of neurogenic lower urinary tract dysfunction, but none can restore normal function. On the other hand, novel neuromodulatory approaches such as epidural spinal cord stimulation (ESCS) of the lumbosacral spinal cord have shown potential to activate neural networks associated with bladder function in rodents with SCI and thus lead to a degree of functional recovery^40,41^. Additionally, clinical studies have shown that transcutaneous electrical spinal cord stimulation (TSCS) – i.e., a non-invasive neuromodulation therapy that stimulates the spinal cord from the surface of the skin – can reengage the spinal circuits involved in bladder control and normalize bladder and urethral sphincter function in patients with SCI^42,43^. Although neuromodulation therapies offer a great promise for restoring normal lower tract function, their mechanism of action (MOA) remain elusive. This is mainly due to the lack of a monitoring modality that can characterize the effects of neuromodulation on spinal cord activity. Combining fUSI with neuromodulation of spinal networks has considerable potential in gaining a better understanding of the MOA of neuromodulation and augmenting its efficacy in improving bladder control in patients with neurogenic lower urinary tract dysfunction. Fine-tuning stimulation wave properties, such as amplitude, frequency, and shape, using fUSI has the potential to facilitate the objective identification of efficacious targets for neuromodulation.

### Limitations and Clinical challenges

While fUSI is a novel technology that enables the monitoring of brain and spinal cord activity, the skull and the lamina bone attenuate and result in aberrant acoustic waves at high frequencies, substantially reducing signal sensitivity. For this reason, most fUSI applications are minimally invasive – with few exceptions such as in young mice (8-12 weeks old with thin skull)^44^ and in pediatric transfontanelle-imaging^17,45^. Surgical procedures to produce a craniotomy^21^ or thinned-skull window^46^ in brain research and laminectomy^26,30^ in spinal cord research are required to harness the host of fUSI benefits. Hence, monitoring spinal cord neuroactivation with fUSI in awake adults is challenging and has yet to be proven. However, recent studies in brain research provide evidence that non-invasive fUSI is capable either through a permanent “acoustic window” installed as part of a skull replacement procedure following a decompressive hemicraniectomy (partial skull removal)^47^ or by intravenously injecting microbubbles-contrast agents for enhancing the fUSI signal^48,49^. Although these approaches have not yet been tested in spinal cord research, the promise of fully noninvasive fUSI in spinal cord is imminent.

It is important to acknowledge that we recorded activity in the thoracic cord (T10 lamina), although the main control mechanism of the bladder is thought to be located in the sacral cord between S2-S4, with the major portion at S3^2,50^. This is a typical limitation in clinical studies that often we are not able to record activity directly from the desirable locations. In our study, we image the spinal cord during urodynamically-controlled micturition in patients undergoing ESCS surgery for chronic low back pain treatment. The midline of the spinal cord at the T10 lamina is the preferred location for insertion of a more rostral spinal cord stimulator and therefore the laminectomy allows us to perform functional neuroimaging only at the T10 region. However, this clinical limitation does not affect our main finding that the hemodynamic signal within the T10 area encodes bladder pressure. In fact, this finding supports the prevailing hypothesis that micturition is regulated by neural circuits that traverse the entire central nervous system from the sacral cord to the prefrontal cortex and vice versa. When the sacral cord receives the sensory information from the bladder, this signal travels up the spinal cord to higher centers in the pons and above^12^. Also, the signal from the brain in turn travels down to the spinal cord to make sure that we only urinate when and where is appropriate ^51^. Therefore, it is likely that the bladder pressure-related signal that we detect at the T10 vertebral body level is a combination of the signal initiated at the sacral cord that traveled towards higher brain centers, and the signal that is transferred from the brain to the bladder through the spinal cord.

Furthermore, while it is common in animal spinal cord studies to perform large laminectomies, retract back muscles and remove connective tissues^26–29^, it is not possible to modify the surgical protocol in order to improve the quality of the fUSI images in human experiments. Instead, we performed only partial and small laminectomies to avoid spine destabilization. In particular, the width of the laminar opening (about 11 mm) was smaller than the width of the ultrasound probe (12.8 mm) and consequently the probe did not perfectly abut the dura. Therefore, it is challenging to image the exactly same 2D plane across patients. Although the imaging planes vary slightly across the 4 patients, this does not affect the spatiotemporal pattern of the hemodynamic signal in the bladder pressure-related regions. In fact, this highlights the strength and robustness of fUSI to overcome the potential to image different 2D slices of the spinal cord across patients.

## Conclusions

Taken together, we present the first in-human characterization of spinal cord hemodynamics during physiological activation of the bladder. By combining fUSI with urodynamically-controlled bladder filling and emptying in human patients with spinal cord laminectomy, we identified spinal cord regions where the hemodynamic signal is strongly correlated with the bladder pressure. These findings demonstrate the existence of a network that is involved in micturition, and open new doors for further investigation of neural network circuits that control and regulate other body processes in healthy and disease conditions.

## Materials and Methods

### Patient and surgical procedures

A total of four participants were imaged continuously during bladder filling and emptying in this study. The participants were recruited from patients who underwent standard-of-care implantation of a spinal cord stimulator paddle lead (PentaTM model 3228) at the Keck School of Medicine of the University of Southern California (USC). All patients were diagnosed with failed back surgery syndrome, which required a T10 partial laminectomy for insertion of stimulation paddle lead in the prone position under general anesthesia.

Spinal cord hemodynamic responses to bladder filling and emptying were acquired via insertion of a fUSI probe into the T10 partial lamina opening prior to placement of the paddle lead (Fig. 1A). Informed consent was obtained from all patients after the nature of the study and possible risks were clearly explained, in compliance with protocols and experimental procedures approved by the USC Institutional Review Board.

### Patient bladder pressure signal acquisition

The urodynamic assessments in this study were conducted using the Laborie Goby (TM) urodynamics system to fill, empty and acquire continuous intravesical bladder pressure measurements of patients. A LaborieT-DOC (TM) catheter was inserted into the bladder, after patients were anesthetized. The position was confirmed by irrigation and aspiration. The infusion port of the catheter was connected to a drainage bag and the manometer port was connected to the Laborie UDS Roam Bluetooth transmitter. The patients were then positioned prone. To begin experiments, the infusion port of the catheter was connected to the infusion tubing and fUSI recordings were performed simultaneously with the urodynamics (See details of the experimental protocols below).

### Functional ultrasound imaging data acquisition

The spinal cord hemodynamic signals were acquired with a fully featured commercial Iconeus One (Iconeus, Paris, France) fUSI system. A 128-element linear array transducer probe with a 15 MHz center frequency and 0.1 mm pitch was inserted through the laminar opening to generate fUSI images (Fig. 1A). This approach enables image acquisition with spatial resolution of 100 μm × 100 μm in-plane, slice thickness of 400 μm, and FOV of 12.8 mm (width) × 10 (depth) mm. The penetration depth was sufficient to image the dorsal portion and part of the ventral portion of the spinal cord on a transverse orientation. The probe was fixed steadily throughout experiments with the FOV transverse and intersecting the spinal cord central canal (Fig. 2). Each image was obtained from 200 compounded frames using 11 tilted plane waves separated by 2° (i.e., from -10° to +10° increment by 2°), at a 500 Hz frame rate. Imaging sessions were performed using a real-time continuous acquisition of successive blocks of 400 ms (with 600 ms pause between pulses) of compounded plane wave images, with a 5500 Hz pulse repetition frequency (PRF). The acoustic amplitudes and intensities of the fUSI sequence remained below the Food and Drug Administration limits for ultrasonic diagnostic imaging (FDA, 510k, Trace 3).

### Experimental protocol

A 26-min continuous fUSI signal acquisition protocol was employed for all patients. The protocol consisted of 5 min fUSI spinal cord baseline recording followed by simultaneous bladder intravesical pressure signal and fUSI signal acquisition, including 2 bladder filling cycles and 1 emptying cycle, interspersed by 2 hold periods and a wash-out period at the end (Fig. 1B). At the 5-min mark, we filled the patients’ bladder through a catheter with 600 ml of saline at a rate of 90 ml/min for approximately 6 min and 40 s, while simultaneously recording the bladder pressure. The filling was paused for about 1 min and 30 s, followed by additional bladder filling with saline for about 1 min. We then stopped the pump for 1 min and 30 s and reversed the pump to continuously withdraw saline via the catheter for 7 min and 40 s at a rate of 90 ml/min, with continuous recording of the bladder pressure. The pump was turned off, then followed by approximately 2 min and 20 s of additional fUSI spinal cord and bladder pressure signal recordings.

## Data analysis

### Data preprocessing

A built-in phase-correlation based sub-pixel motion registration^52^ and singular-value-decomposition (SVD) based clutter filtering algorithms^53^, in the Iconeus One acquisition system were used to separate tissue motion signal from blood signal to generate relative pD signal intensity images^54^. We adopted rigid motion correction techniques^55^ that have successfully been used in fUSI^21,23,30^ and other neuroimaging studies^56–58^, to address potential physiological and motion artifacts unique to human spinal cord imaging. This was combined with in-house high frequency smoothing filtering. We utilized a lowpass filter with normalized passband frequency of 0.04 Hz, with a stopband attenuation of 60 dB that compensates for delay introduced by the filter, to remove high-frequency fluctuations in the pD signals.

### Spatiotemporal correlation of bladder pressure changes to ΔSCBV

We assessed the spatiotemporal effects of bladder filling and emptying on spinal cord hemodynamics. We generated pixel-wise activation time course curves of ΔSCBV as a percentage change of the pD signal relative to baseline activity for the whole spinal cord FOV. The mean pD signal activity acquired 1 min preceding the onset of the bladder filling was utilized as the baseline for the analysis. We investigated whether there are spinal cord regions where ΔSCBV is correlated with the bladder pressure during filling and emptying. To test this hypothesis, we computed Pearson correlation coefficients for each pixel in the spinal cord fUSI image. To this end, the %pD signal intensity time series curve from each pixel is correlated with the bladder pressure signal across time to determine pixels with statistically significant correlation (p < 0.01, with FDR correction). We generated statistical correlation activation maps of the pixels that show significant positive and negative correlations above an r-coefficient threshold (r > 0.35 and r < -0.35). Finally, to visualize the temporal dynamics of the percentage ΔSCVB, we derived the mean % pD signal change curves from averaging the signal over the pixels with significant correlation to the bladder pressure signal.

### Decoding bladder pressure dynamics from SCBV signals

Next, we attempted to identify spinal cord regions with ΔSCBV that captures the temporal changes of the bladder pressure, without direct knowledge of the bladder pressure signal. We utilized a machine learning algorithm cPCA+SVM that includes the following steps: 1) align the preprocessed SCBV signals extracted from the bladder filling and emptying time epochs, 2) reduce data dimensionality and select features that optimally allow discrimination between filling and emptying states, 3) dissociate and identify relevant spinal cord areas that encode the bladder pressure dynamics and 4) cross validate and evaluate the decoder performance (Fig. 4). To do so, we determined the percentage change in pD signal in each pixel of the fUSI images extracted during the filling and emptying epochs for each patient, relative to reference signal activity. The signals acquired 30 s just before the onset of filling and 30 s before onset of emptying (i.e., during the 2^nd^ hold period) were used as reference to calculate the %pD signal change for each pixel during the filling and emptying periods, respectively. The entire fUSI spinal cord 2D image space was utilized in the machine learning algorithm (Fig. 4A). Each 2D time series data was vectorized to 1D vectors and aligned in rows to form 2D (pixels × time) matrix classes (filling – class:0 and emptying – class:1) (Fig. 4B). We employed classwise principal component analysis (cPCA) ^31,32^ and support vector machines discrimination (SVM) ^59^, to reduce data sparsity and dimensionality while maintaining enough components to retain over 95% variance in the data and to select the most relevant subspaces to separate the classes. SVMs provide a set of supervised learning tools for classification that are effective for high-dimensional spaces even when the feature dimensions are larger than the number of samples – such as the data employed in this study. We combined cPCA with SVM to classify the cPCA-transformed fUSI image into filling (class:0) and emptying (class:1) bladder pressure states. This analysis provides weights that reflect the most relevant pixels used for classifying between classes (Fig. 4C). The relevant pixels represent a feature extraction mapping to the 2D spinal cord image space and are derived from the two 1D low-dimension subspaces that are optimized for each class. The subspaces identify pixels in the spinal cord fUSI images that encode the differences between the filling and emptying classes, when projected back to the image space. Each pixel is assigned a relative weight of relevance (normalized between [-1 1] – pixels with values close to +1 or -1 indicate high relevance components, while pixels with values close to 0 are less important and whose fluctuations are likely due to noise). ΔSCBV of pixels with positive and negative relative weights of relevance are positively and negatively correlated with bladder pressure changes, respectively.

### Optimal data amount for training and cross-validating the classifier

Next, to investigate the optimal amount of data needed to generate the best correlation between ΔSCBV and bladder pressure, we employed a similar cPCA+SVM analysis as outlined above with a sliding window of cumulative data amounts. We utilized 30 s of initial data followed by 10 s increments to derive the cumulative data used to train and cross-validate the classifier. We assumed that data acquired at the end of the filling period are more comparable to the data acquired at the onset of emptying and thus, we accumulated the filling data in reverse order (Fig. 4D). We followed comparable cPCA and SVM classification steps as outlined above with increasing data amounts. Each data amount produced a corresponding relevant weighted pixels-matrix and associated mean % pD signal change time course curves (Fig. 4D, highlighted panel), relative to the reference activity. To determine the optimal amount of data and pixel weights, we utilized the mean % pD signal changes derived from the weighted regions for each patient to determine the correlation between the pD signal curve resulting for each cumulative data amount and the bladder pressure signal (Fig. 4D, highlighted panel). The data amount corresponding to the highest correlation coefficient was utilized to select the optimal pixel weights and % pD signal change curve. To cross-validate the classification analysis, we allocated a subset from each data class for training (80%) and testing (20%).

### Software analysis

All data pre- and post-processing and statistical analysis were performed using Matlab Version 9.13.0.2193358 (R2022b).

## Data availability

The datasets generated and analyzed during the current study are available from the corresponding authors on reasonable request.

## Acknowledgments

We thank the participants that made this study possible. This work was supported by “The USC Neurorestoration Center” at the University of Southern California, “The Hellman Foundation” and the “Marlan and Rosemary Bourns College of Engineering” at the University of California Riverside through start-up funding.

## Author Contributions

D.J.L., E.I.K, V.R.E., C.L. and V.N.C conceived the study.; D.J.L. performed the surgeries; D.J.L., W.C., A.A., Y.T.L., and K.W. acquired the functional ultrasound data; K.A.A. performed the functional ultrasound data processing, statistical and machine learning decoding analysis; K.A.A., D.J.L. and V.N.C drafted the manuscript with substantial contribution from J.R., E.I.K, V.R.E. and C.L.; All authors edited and approved the final version of the manuscript; C.L. and V.N.C. supervised the research.

